# The flower pot method of REM sleep deprivation causes apoptotic cell death in the hepatocytes of rat

**DOI:** 10.1101/375717

**Authors:** Atul Pandey, Devesh Kumar, Gopesh Ray, Santosh Kar

**Author notes:** **Corresponding Authors** &.

## Abstract

**Introduction:** The rapid eye movement sleep deprivation (REMSD) of rats relates with increased inflammations, acute phase response, oxidative damage, neuronal cell loss, and neurodegenerative diseases. Whereas, its role outside brain are not well studied. This study tried to explore the causal effect of REM sleep loss on hepatocytes.

**Methods:** We deprived the rats of REM sleep using standard flower pot method. We focused on liver to see the REMSD affects which controls most of the metabolic processes of the body.

**Results:** We report here that flower pot induced REMSD causes apoptotic cell death of hepatocytes (~10% by Annexin Assay & ~20% by TUNEL assay). This were further got alleviated up to extent after sleep recovery of 5 days (recovered approximately 8.0% by Annexin Assay & 14% by TUNEL assay). The gene expression and protein level profiling revealed the up-regulation of p53, Bax, Cytochrome c, Caspase 3, and Caspase 9. While, Bcl2 which is an anti-apoptotic protein were down-regulated in response to REMSD. Relentless recovery of 5 days affected the expression pattern of these genes/proteins.

**Conclusions:** Our study offer great pathological and physiological significance for sleep loss, by inferring the apoptotic cell-death in the hepatocytes of rat. This further signifies the functional and preventive role of REM sleep which is unique to mammals and avians with certain exceptions, as its loss can affect the natural well-being and survival of the individuals.

**Highlights of the study:**

- We observed significant apoptosis in the hepatocytes of REMSD group of rats.
- Our expression analysis confirmed altered expression for genes p53, Bcl2, Bax, and Caspase-3 after REMSD.
- Protein level analysis supported our gene expression results for p53, Bcl2, Bax, Caspase 3 and Caspase 9 after REMSD.
- Sleep recovery improved the respective genes and protein expression levels towards normalcy, signifying the functional role of REM sleep.

## 1.0. Introduction

Sleep is an important evolutionary physiological and behavioral process required for the survival and well-being of animals studied, although no predominant hypothesis has emerged to explain its functions [1,2]. In mammals, it is categorize as two types, non-rapid eye movement (NREM) sleep and rapid eye movement (REM) sleep. Evolutionary REM sleep is present in mammals and birds with essential functions related to physiological and ecological success. REM sleep, also called as paradoxical sleep in general associates with memory consolidation, brain maturation, spatial memory acquisition, muscle regeneration and maintenance of body physiology [3–9]. Functional aspect of REM sleep can inferred by its effect on amygdala activity in response to previous emotional experiences, reorganizing the hippocampal excitability, pruning and maintenance of new synapses in development and learning [10–12]. Its prolonged loss can alter blood brain barrier functions and can be fatal[13–15]. Some recent studies have shown that REMSD can cause apoptosis in rat’s brain [16–18]. Our previous reports, we had shown that REMSD induces acute phase response in the liver, increasing the synthesis of pro-inflammatory cytokines like IL1β, IL-6 and IL-12 and increasing the levels of liver toxicity marker enzymes, alanine transaminase (ALT) and aspartic transaminase (AST) which circulates in the blood [19]. Whereas, we also found that REMSD can induce the ROS level and NO production in the hepatocytes along with making them more susceptible to oxidative stress [20]. Recent, reports also suggest the effect of REMSD on the new born baby’s sleep [21]. Liver being the metabolic hub contributes to the maintenance of body physiology and well-being. It synthesizes complement proteins and houses Kupffer cells and natural killer (NK) cells, which are important components of the innate immune system. We hypothesized here based on our previous reports and literature that REMSD will might affect the survival of liver cells.

Apoptosis, a regulated form of cell death with many check points and molecular mediators, can initiate in hepatocytes via the death receptors mediated extrinsic pathway, or by cellular perturbations that constitute the intrinsic pathway [22]. Unlike other cells, in hepatocytes, both the pathways converge on mitochondria. The mitochondrial permeabilization is enough to induce apoptosis of hepatocytes [23,24]. Generally, apoptosis can aggravate tissue injury, inflammation and fibrosis [25–27]. Apoptosis of hepatocytes were also evident in viral hepatitis, metabolic diseases, alcoholic steatohepatitis, autoimmune hepatitis and drug induced liver injury which links liver injury to death of hepatocytes [28–32]. In current study, we observed that considerable REMSD in rats induced apoptosis of hepatocytes by 4^th^ & 9^th^ day as indicated by DNA laddering, Annexin V labeling and Terminal deoxynucleotidyl transferase dUTP nick end labeling (TUNEL) assays. Furthermore, real time PCR and western blot analysis revealed the downregulation & upregulation of anti-apoptotic and pro-apoptotic genes like Bcl2, Bax, Caspase 3 and p53 in the hepatocytes of REMSD rats. Sleep recovery affected the expression levels of all genes/proteins studied indicating its redressal effects. Our results reveal that REM sleep is an important phenomenon for individual well-being and its loss can induce the cell death in hepatocytes. These findings further support the protective and adaptive evolutionary role of the REM sleep for the maintenance and survival of the organisms.

## 2.0. Materials and Methods

We used male wistar rat’s weighing 220-260gms for study. We kept rats in the institutional animal house facility of the University with a 12:12hrs L:D cycle (lights on at 7.00 am). We supplied food and water *ad libtium* for all experimental groups during treatments. We got approval from University’s Institutional Animal Ethics Committee for all protocols.

### 2.1. Methods used for REM sleep deprivation and recovery

We used classical flower-pot method for depriving the REM sleep in rats [33,34]. We kept rats for REM sleep deprivation on small raised platform (6.5 cm in diameter) surrounded by water. We maintained the large platform control (LPC) group of rats on platform of 12.5cm diameter. Meanwhile, the cage control (CC) rats remained in cages during experiments. In REMSD groups, rats only could sit, crouch and have non-REM sleep, while no REM sleep. The muscle atonia associated with REM sleep stage forced them to awake and thus deprived of it. We allowed the rats to have 5 days of sleep recovery (for recovery group) after 9 days of REMSD. We terminated the experiments after 4 days, 9 days and 5 days of sleep recovery. We collected the tissue/cells from individual rat and did further analysis.

### 2.2. Histopathological examination

We anesthetized the rats, perfused their liver and dissected them out. We fixed the liver after collection for 3 days in 4% para formaldehyde-phosphate buffered saline (0.01 mol l-1 phosphate buffer, 0.0027 mol l-1 KCl, 0.137 mol l-1 NaCl, pH 7.4) solution. We embedded the tissue into paraffin, made 5-6 mm thick sections and mounted on the glass microscope slides using standard histopathological techniques. The sections were stained with hematoxylin & eosin (H&E) and examined by light microscopy [35]. The liver from cage control and LPC treatment group rats were also processed like experimental groups. The pathologist who examined the slides was blind to the treatments. We marked the inflammatory cells detected in examinations on slides.

### 2.3. Hepatocytes preparation

We isolated the hepatocytes from individual liver of CC, LPC, and REMSD group of rats [36]. In brief, we opened the abdomens of rats through a mid-line incision. We placed the portal cannula inside the liver and perfused it with 0.02% EDTA sloution at 37° C. The flow rate was 30 ml per minutes and perfusion took on average 15 minutes. We recirculated the collagenase solution (37° C) through cannula at same flow for 15 minute. We disrupted the liver capsules after perfusion and digestion. The parenchyma cells were suspended in the ice-cold Hank’s balance salt solution. We washed the cells by centrifuging them at 500 rpm for 5 min 2-3 times. We further centrifuged cells over 30% percoll at 100g for 5 min to increase the purity. Viability at the time of labeling as measured by Trypan blue exclusion was ≥ 95%.

### 2.4. Annexin V labeling of hepatocytes for detection of Apoptosis

We performed Annexin V assay using the Flow-TACS kit (4835-01-K) from R&D systems. We analyzed the labeled samples by flow cytometer (BD FACS Calibur) within an hour. We considered Annexin V+ and PI-cells as early apoptotic. Annexin V +and PI+ cells as late apoptotic whereas Annexin V-& PI+ cells to be necrotic.

### 2.5. TUNEL labeling of hepatocytes DNA for detection of Apoptosis

We performed TUNEL assay following instructions from Flow-TACS kit (R&D systems, 4817-60-K). We analyzed the samples by flow cytometer (BD FACS Calibur) within an hour. We considered TUNEL+ and PI-cells as early apoptotic. While, TUNEL+ and PI+ cells as late apoptotic and TUNEL- and PI+ cells as necrotic cells.

### 2.6. Isolation of RNA and TaqMan Real-time PCR

We isolated total RNA using RNA purification kit (Qiagen) from hepatocytes stored in RNA later (Sigma, R0901). We assessed the RNA concentration and integrity by Nanodrop and Agilent 2100 Bioanalyzer (Agilent Technologies, Massy, France). We prepared cDNA from total RNA using the reverse transcription PCR kit (Applied Biosystems). The GADPH served as a housekeeping gene and CC group as calibrator probes. The reporter dye were FAM labeled on 5’ end and quencher VIC labeled on 3’ end. We obtained PCR master mix, and PCR plate/tubes from Applied Biosystems. We followed manufacturer’s instructions of respective kits. The catalogue number of gene probes were GAPDH (Rn01749022_g1), Bcl2 (Rn99999125_m1), Bax (Rn02532082_g1), p53 (Rn00755717_m1), Caspase 3 (Rn01488068_g1), and master mix (Rn99999916-g1).

### 2.7. Western Blot Analysis

We performed western blot analysis of our protein samples as described [37]. We used primary antibody in 1/1000 dilutions (p53-sc126; Bcl2-sc23960, Bax-sc7480, caspase 3-sc136219; caspase 9-sc81650; cytochrome-c-sc13156; GAPDH-sc365062, Santacruz Biotechnology, Inc, USA). We used horseradish peroxidase-conjugated (sc-516102) secondary antibody (1/5000 dilutions, Santacruz Biotechnology, Inc, USA). We developed membranes using ECL reagents and acquired images (Photo and Imaging 2.0; Hewlett-Packard). We used Adobe (Photoshop 8.0) software for image analysis.

### 2.8. Statistical Analysis

We used Graph pad-Prism 5 (version 5.01) for Tukey’s HSD posthoc test following ANOVA for measuring out the effect across treatment groups. We considered p values < 0.05 significant for our analysis. We used R^2^ values to show the size effects.

## 3.0. Results and Discussions

There are inconsistent reports of sleep loss and cellular apoptotic responses. Some studies reported no evidence of brain-cell degeneration after sleep deprivation in rats [38,39]. Yet, recent few studies recorded apoptotic neuronal cell degeneration against sleep deprivation [16,18,40–43]. Here, we investigated the effect of REM sleep loss and effects on liver. We examined the liver sections of rats after hematoxylin/eosin (H&E) staining on different days of REMSD and after 5 days of recovery (5DR) and compared them with control groups. The CC group and LPC group of rats showed normal histology on 4th day (Fig.1a & b) as well as on 9th day (Fig.1d & e) of experiment. While, sections of REMSD group rats on 4th day showed mild lymphocytic infiltration, dilated central vein and parenchymatous cells injury (Fig.1c). The sections of 9th day showed hepatic degeneration, more lymphocytic infiltration (Fig.1f). Sleep recovery of 5 days improved the situation and showed mild lymphocytic infiltration (Fig.1g). A previous study supports our finding, where total and partial sleep deprivation affected liver, lungs and small intestine of rats causing oxidative DNA damage [44]. Hepatic steatosis, and mucolipidosis were detected in the liver of REM sleep deprived rats after 1, 3 and 5 days of deprivation [45]. Previously, prolonged sleep deprivation was found associated with disturbed liver functions, and hyperphosphatemia involving human volunteers [46].

**Figure 1:**
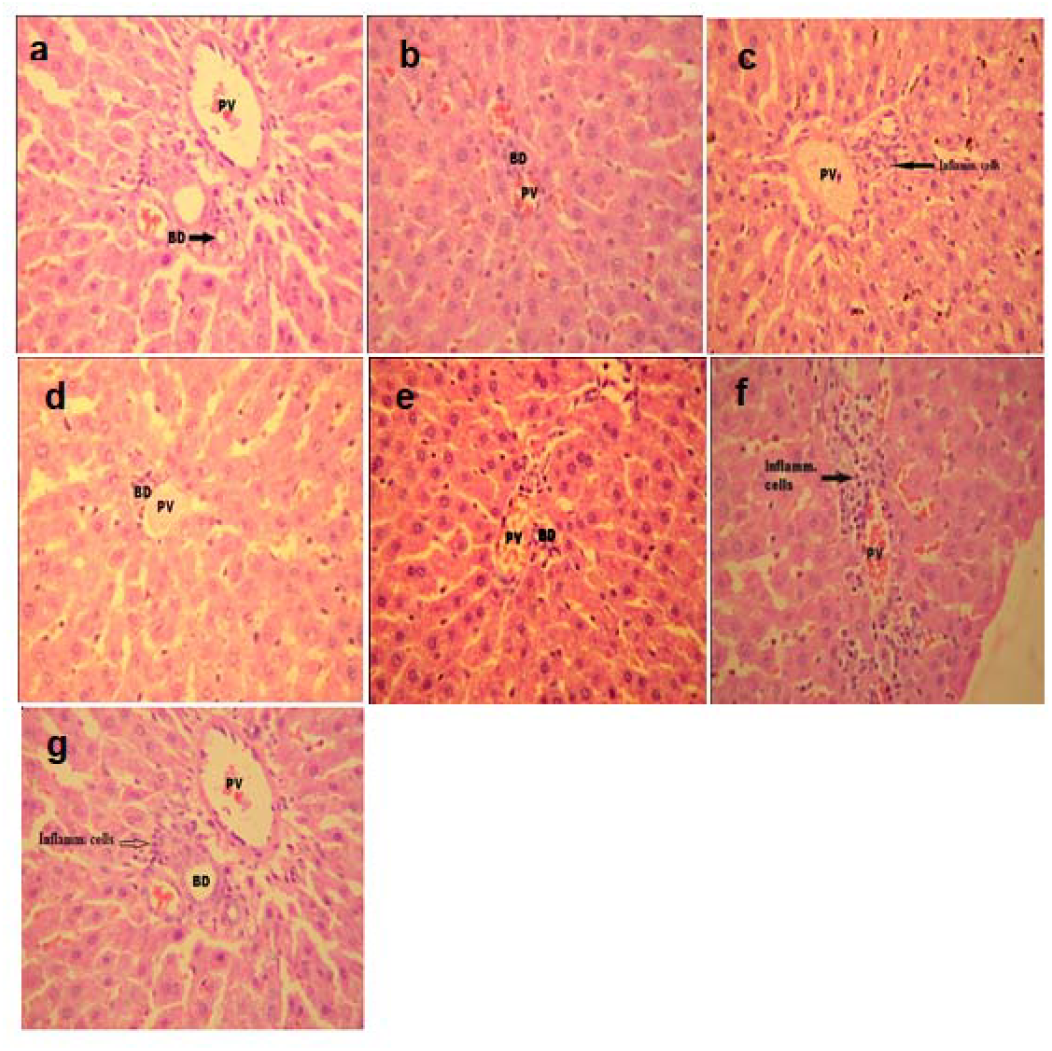
Histopathological analysis of rat liver tissue from cage control, large platform control and REM sleep deprived group. Hematoxylin/eosin (H&E) stained sections of the liver in progression with REMSD in rats showing inflammatory cells. (Original magnification, 200x). The sections show portal vein (PV) as well as bile duct (BD) along with hepatocytes, kupffer cells and sinusoids. (a) CC 4^th^ day, (b) LPC 4^th^ day, (c) REM sleep deprived 4^th^ day, (d) CC 9^th^ day, (e) LPC 9^th^ day, (f) REM sleep deprived 9^th^ day and (g) REM sleep deprived 5 day recovery group.

Based on our histopathology results, we hypothesized and tested that REMSD might will cause cell death in hepatocytes. Our hypothesis was further supported by report of cell death in the peripheral organs and tissues due to total and selective sleep loss in rats [44], and our recent reports suggesting REMSD induced increased acute phase response in serum and ROS in hepatocytes [20,47]. We labeled hepatocytes with Annexin V and TUNEL to estimate the cell death. We reported increased number of Annexin V positive hepatocytes in the REMSD group by 4^th^ day as well as by 9^th^ day of sleep deprivation compared to CC and LPC control groups, while, sleep recovery of 5 days decreased the number of Annexin V positive cells (Fig. 2A, One way ANOVA F=84.13, df=6, p<0.001). Further, we stained hepatocytes for TUNEL-positive nuclei after 4^th^ and 9^th^ days of REMSD and 5DR and observed the significant change in labeling. We observed quite significant number of cells positive for TUNEL by 4^th^ (10±2.75%) and 9th (20.54±2.02%) day of experiment in comparison to controls (Fig. 2B, One way ANOVA F=181.72, df=6, p<0.001). The LPC group of rats didn’t show significantly more labeling for Annexin V and TUNEL compared to cage control, which indicated that stress due to confinement is not contributing for this (Fig. 2B). TUNEL assay is most often used to identify apoptosis of cells [48]; however, recent reports have indicated that TUNEL can detect DNA fragmentation in necrotic tissues [49]. The non-specific staining in TUNEL assay was analyzed by using hepatocytes from CC and LPC groups of rats. Both the control groups showed small percentage of cells (<2%) being stained in TUNEL assay indicating that there was no major nonspecific staining.

**Figure 2:**
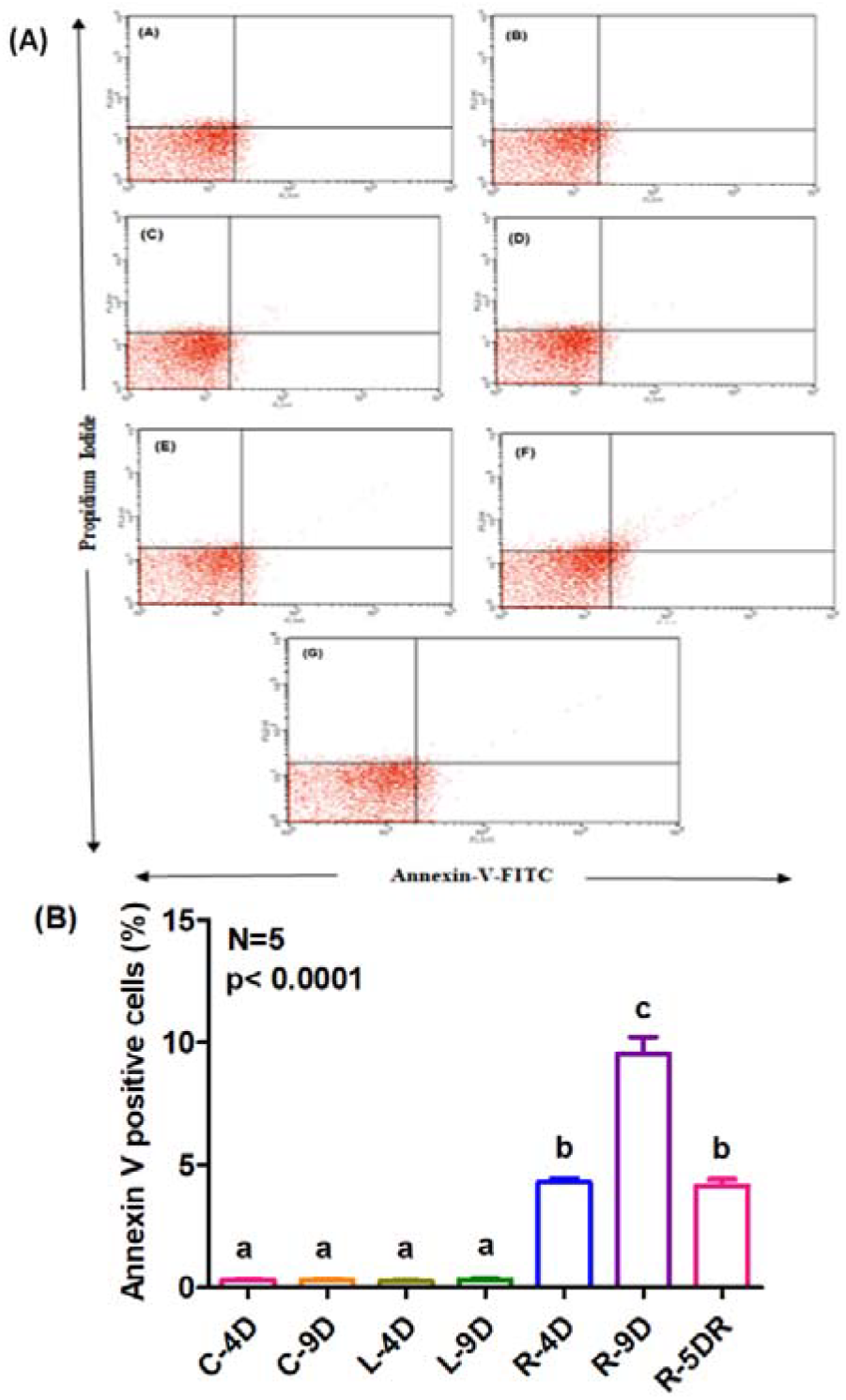
Detection of Annexin V positive cells in the hepatocytes of rats; (A), Annexin V labeling of hepatocytes for CC (Aa-4day &Ab-9day), LPC (Ac-4day &Ad-9day) and REM sleep deprived group of rats (Ae-4day, Af-9day & Ag-5day recovery). X-axis represents the labeling for Annexin V-FITC while Y-axis represents the labeling of Propidium iodide. (B), Percentage average labeling of Annexin V-FITC labeling of hepatocytes for CC, LPC and REMSD group of rats. X-axis represents the treatment groups and Y-axis represents the average percentage of Annexin V positive cells. Treatment groups marked with different small letters are statistically different in a Tukey post-hoc tests followed by ANOVA. P value < 0.05 was considered statistically significant. [CC-4=Cage control-Day 4, LPC-4=Large platform control-Day 4, CC-9= Cage control-Day 9, LPC-9=Large platform control-Day 9, REM-4=REMSD-Day4, REM-9=REMSD-Day 9 and REM-5DR, 5 days of sleep recovery after 9 days of sleep deprivation]

The above results indicates that flower plot induced sleep deprivation can cause hepatic cell death with unknown reasons. To elucidate this further, we looked into literature and tried to correlate our findings with known knowledge. A previous studies suggest that even moderate levels of continuous stress can induce apoptosis in cultured hepatic cells [50–53]. In our experimental model system, we can’t differentiate out between experimental born stress and loss of REM sleep. We considered LPC group of rats as our sham control group which were also subjected with similar stress and isolation. So, we moved on with this notion of comparing LPC group from control for non-specific stress related effects. In future, identification of relevant gene and involvement of gene surgery (CRISPR-Cas) like technology will resolve this issue of taking out the sleep deprivation procedure related stress. Thus, will help in more clear way to understand the consequence of REM/total sleep deprivation.

The production of superoxide’s due to oxidative stresses can kill hepatocytes involving apoptosis. This includes activation of caspases and the c-Jun N-terminal kinases [54]. In general, the phenomenon of apoptosis at cellular levels are highly correlated with iNOS induction and concomitant massive and sustained circulation of NO. Whereas, sustained NO circulation is strongly correlated with high ROS levels and can be observed by chromatin condensation and DNA laddering. While, tumor suppressor p53 precedes DNA fragmentation in cells in response to NO generation [55]. Studies further suggest that cytokines like TNF-α, IL-1β, and INF-γ synergistically activate iNOS expression in the liver. While, NO exerts a protective effect both *in vivo* and *in vitro* by blocking TNF-α induced apoptosis and hepatotoxicity, by inhibiting the caspase-3-like protease activity [56,57]. Cellular susceptibility toward NO varies between different types of cells and tissues [58]. NO has been shown to cause accumulation of the nuclear phospho-protein p53 in RAW 264.7 cells [59]. There is suggestive evidence of the role of p53 in apoptotic pathway in response to DNA damage [55]. Our recent finding suggested that REMSD of rats increased iNOS expression. While, this higher NO production in hepatocytes leads the ROS production and augmented susceptibility of hepatocytes towards oxidative stress [60]. This allowed us to hypothesize that might be caspases are involved in the REMSD driven apoptotic process?

In our experimental condition expression level of p53 gene (Fig. 3A, One way ANOVA F=54.31, df=6, p<0.001) and protein (Fig.4A&B, One way ANOVA F=24.92, df=14, p<0.001) was found increased after days of REMSD. The 5 day sleep recovery improved the circulatory p53 level. Accumulation of the tumor suppressor p53 in response to endogenously generated NO correlates with accompanying event of apoptosis [61]. Exposure of NO, generated from an NO donor or from overexpression of inducible-type NOS, results in p53 protein accumulation [62]. In apoptotic condition, ROS production was observed along with increased synthesis and circulation of NO and p53 genes and further found correlated with Bcl2 family proteins [63].

**Figure 3:**
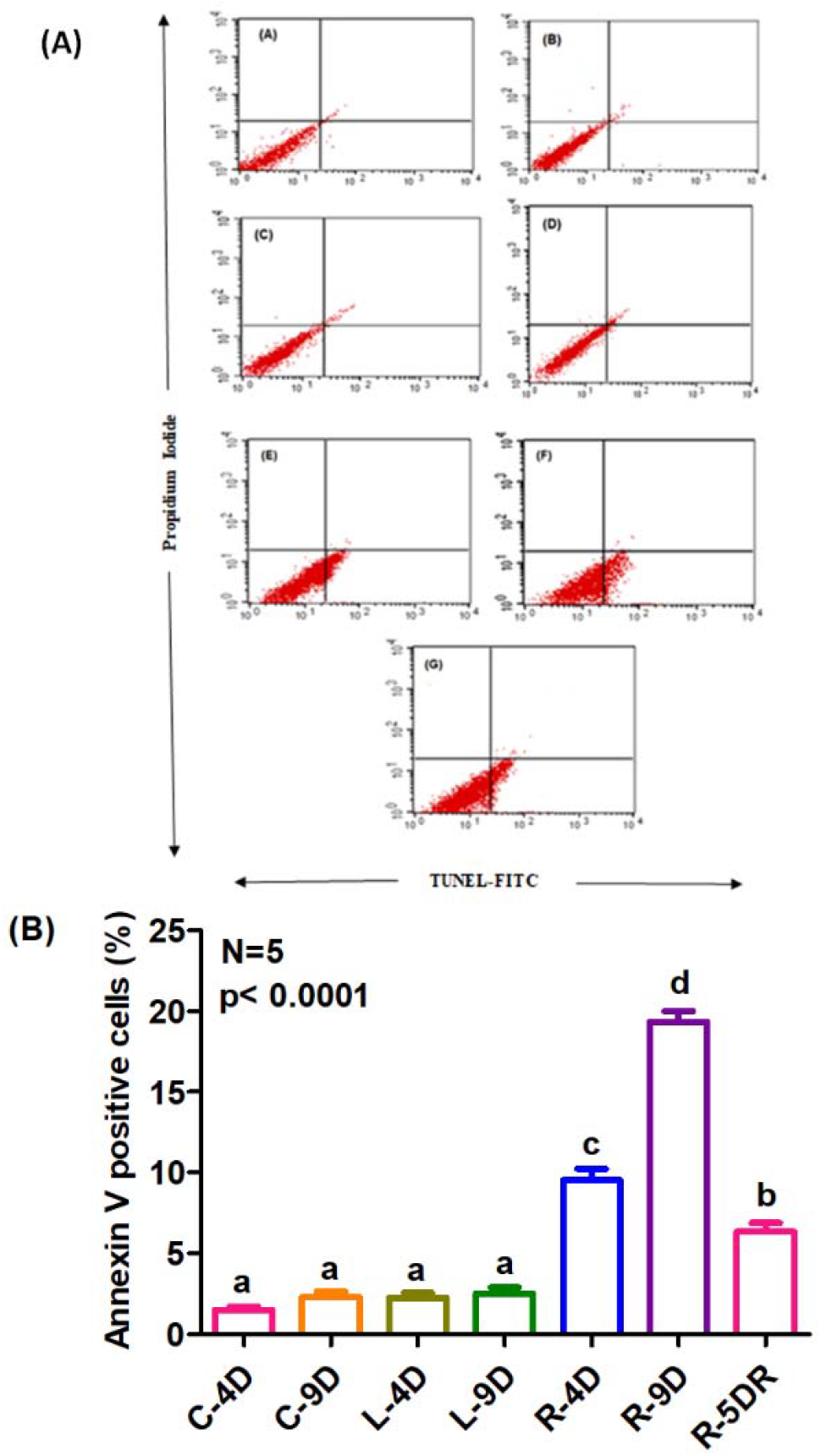
Detection of TUNEL positive cells in the hepatocytes of rats; (A), TUNEL labeling of hepatocytes for CC (Aa-4day &Ab-9day), LPC (Ac-4day &Ad-9day) and REM sleep deprived group of rats (Ae-4day, Af-9day & Ag-5day recovery). X-axis represents the labeling for TUNEL-FITC while Y-axis represents the labeling of Propidium iodide. (B), Percentage average labeling of TUNEL-FITC labeling of hepatocytes for CC, LPC and REMSD group of rats. Other details in figure 2.

**Figure 4:**
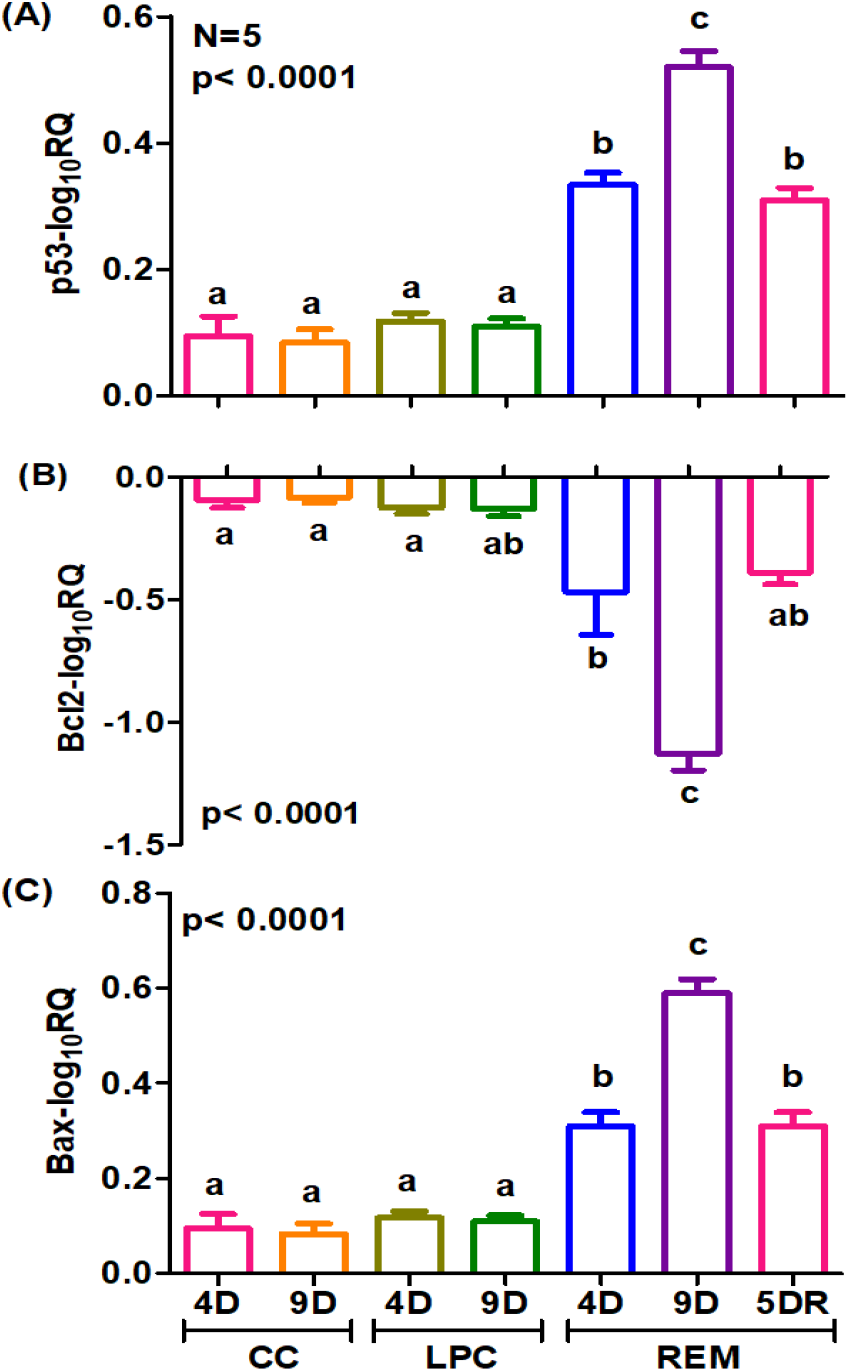
Analysis of apoptotic genes by real time PCR. The graph shows log fold change in expression pattern of p53 genes (a), Bcl2 (b) and Bax (c) respectively. Cage control samples were taken as calibrator while GAPDH was endogenous control for respective genes. X-axis represents the different days of sleep deprivation for treatment groups and Y-axis represents the log fold expression of genes. See figure 2 for further statistical and legend specific details.

Further, we measured the expression of apoptotic genes and proteins in the hepatocytes of rats. The expression pattern of Bcl2, Bax, and Caspase 3 genes were measured and compared between CC, LPC and REMSD group of rats. The expression of Bcl2, an anti-apoptotic gene whose protein product binds to Bcl-x family of pro-apoptotic proteins such as Bax, PUMA, Noxa etc, preventing them to permeabilize the mitochondrial membrane, were found reduced significantly after 4^th^ and 9^th^ day of REMSD (Fig. 2B, One way ANOVA F=24.80, df=6, p<0.001).

We assume that p53 gene which were found over expressed (Fig.4A) due to REMSD and earlier correlated to be involved in apoptosis regulation at an early stage, might have induced the secretion of Bax gene (Fig. 2C, One way ANOVA F=23.89, df=6, p<0.001) and caspase 3 gene (Fig.5A, One way ANOVA F=79.52, df=6, p<0.001). These proteins were supposed to be involved during the final stages of apoptosis [64,65]. We further supported this with caspase 3 protein expression analysis (Fig.5B&C, One way ANOVA F=47.88, df=19, p<0.001).

**Figure 5:**
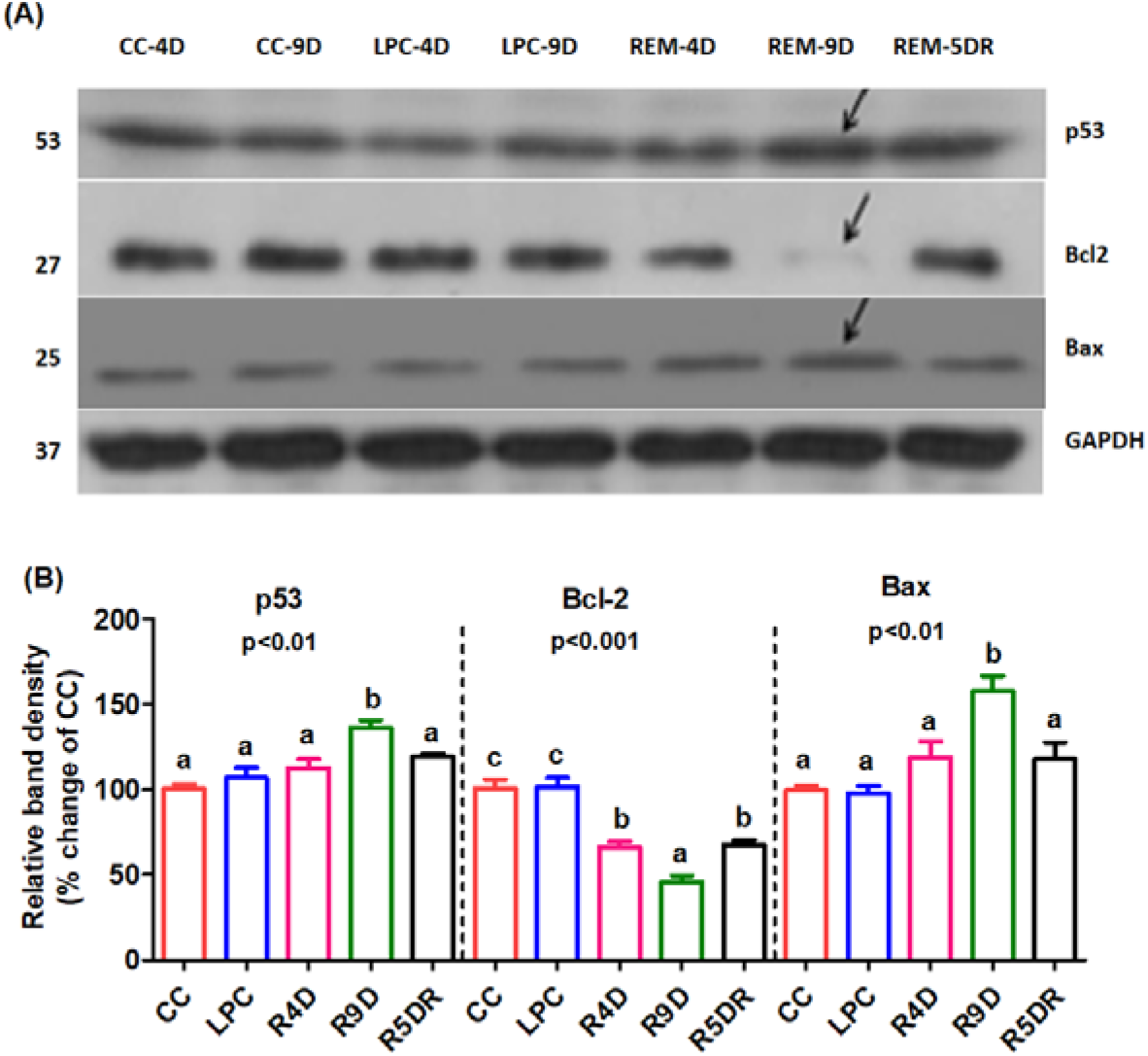
Analysis of hepatocytes proteins using WB from CC, LPC and REM sleep deprived group rats. (a), Lane 1(CC-4D), lane 2 (CC-9D), Lane 3(LPC-4D) and Lane 4 (LPC-9D) represent samples from the CC and LPC group of rats after 4^th^ and 9^th^ day of the start of experiment, while Lane 5 (REM-4D), Lane 6 (REM-9D) and Lane 6 (REM-5DR) represent samples from REMSD group rats after 4^th^day, 9^th^ day and after 5DR of REM sleep deprivation. Glyceraldehyde 3 phosphate dehydrogenase (GAPDH) was used as an endogenous loading control. (b), Densitometric analysis of the protein bands expressed in reference to the endogenous loading control GAPDH. X-axis represents the different days of sleep deprivation for treatment groups and Y-axis represents the gel band density. P value < 0.05 were considered as statistically significant. Other details as in figure 2.

**Figure 6:**
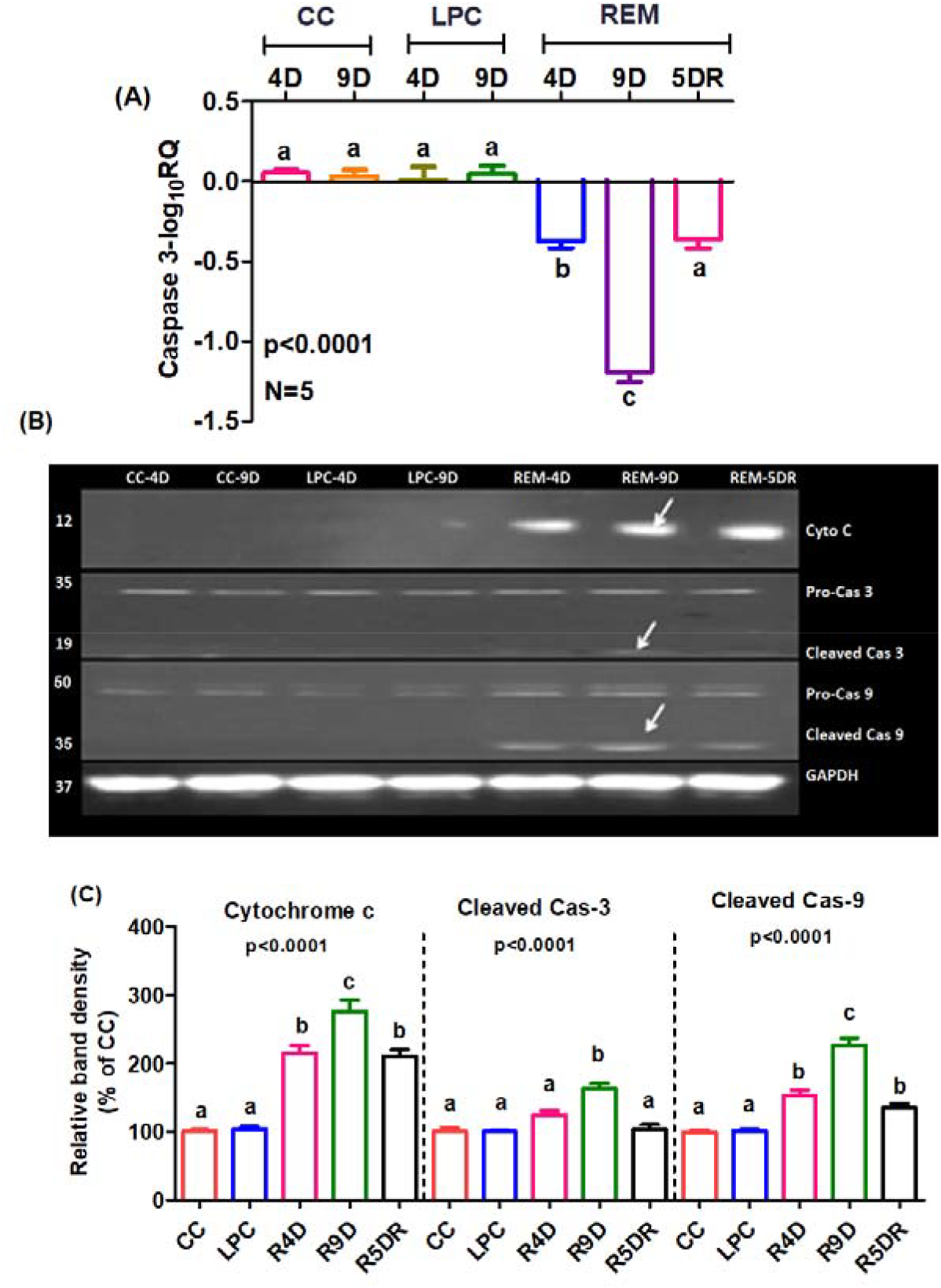
Analysis of Caspases involved in apoptosis from CC, LPC and REM sleep deprived group rats. (A), the graph shows log fold change in expression pattern of caspase 3 gene. Cage control samples were taken as calibrator while GAPDH was endogenous control for respective genes. (B), Lane 1(CC-4D), lane 2 (CC-9D), Lane 3(LPC-4D) and Lane 4 (LPC-9D) represent samples from the CC and LPC group of rats after 4^th^ and 9^th^ day of the start of experiment, while Lane 5 (REM-4D), Lane 6 (REM-9D) and Lane 6 (REM-5DR) represent samples from REMSD group rats after 4^th^day, 9^th^ day and after 5DR of REM sleep deprivation. (C), Densitometric analysis of the protein bands expressed in reference to the endogenous loading control GAPDH. X-axis represents the different days of sleep deprivation for treatment groups and Y-axis represents the gel band density. Value of P < 0.05 were considered as statistically significant. Other details as in figure 2.

Reports suggest that, increase in the level of p53, which also transcriptionally regulates Bax, also contributes the redistribution of Bax from the cytosol to mitochondria [66]. We observed the decreased levels of Bcl2 while increased levels of Bax proteins (Fig. 4A&B). The decreased Bcl2 and increased Bax alters the Bcl2/Bax protein ratio affecting mitochondrial membrane potential. Taken all together these events are reported to facilitate permeabilization of mitochondrial membrane and release of cytochrome C which activate caspases [67–69]. In our experimental conditions also, we observed higher circulation of cytochrome C and activation of Cas 3, and Cas 9 (Fig.5B&C).

We suppose that activated caspases (particularly Cas 3) induced apoptosis of the hepatocytes (Fig.1A&B). In a study reported earlier REMSD of rats was similarly reported to cause apoptosis of neurons involving Bcl2 family proteins and caspases which further supports our current line of observations [16,18]. The physiological consequence of hepatocytic cell death also signifies and justifies the evolutionary and conserved roles of REM sleep which evolved lately in avians and mammals with certain exceptions.

## Acknowledgements

This research was carried out and supported by the lab running grant of the laboratory of SKK at School of Biotechnology, Jawaharlal Nehru University, New Delhi, India. AP was supported by the research fellowship of University grant commission, India, DK was supported by master’s fellowship of department of biotechnology, India, while GK was supported with JRF fellowship of National Medicinal Plant Board, Department of Ayush, India.

## Abbreviations

REM: rapid eye movement
NREM: non rapid eye movement
REMSD: rapid eye movement sleep deprivationn
CC: cage control
LPC: large platform control
TUNEL: Terminal deoxynucleotidyl transferase dUTP nick end labeling
EDTA: Ethylene diamine tetra acetic acid
RIPA: Radio immuno precipitation assay
FITC: Fluorescein isothiocyanate
TDT: Terminal deoxynucleotidyl transferase
FAM: 6-carboxyfluorescein
ROS: Reactive oxygen species
NO: Nitric Oxide

## Compliance with ethical standards

### Funding

This research received no grant from any funding agency in the public, commercial, or not-for-profit sectors.

### Ethical approval

All applicable international, national, and/or institutional guidelines for the care and use of animals were followed.

### Informed consent

Not applicable, subjects involved in study were rodents (Wistar rats).

